# Small Heat Shock Proteins have a Paramount Role in *Trypanosoma cruzi* Infection Impacting Intestinal Homeostasis of an insect vector of Chagas Disease

**DOI:** 10.64898/2026.01.07.698241

**Authors:** Tainan C. Guedes-Silva, Ana B. Walter-Nuno, Jéssica C.T. Pereira, Marianna R. França, Felipe A. Dias, Hugo D. Perdomo, Isabela Ramos, Gabriela O. Paiva-Silva, Rafael D. Mesquita, Pedro L. Oliveira

## Abstract

*Rhodnius prolixus*, an insect vector of Chagas disease, ingests blood meals several-fold its own body weight. This nutritional overload triggers specific adaptive responses that prevent or remediate damage from multiple stressors, including oxidative, osmotic, and microbial challenges. Transcriptomic analysis of first-instar nymph guts revealed five members of the Small Heat-Shock Protein (sHSP) family that were highly expressed after blood meals but were downregulated upon *Trypanosoma cruzi* infection. sHSPs are proteins involved in cellular homeostasis and stress responses. Simultaneous knockdown of all five sHSPs profoundly disrupted insect physiology, causing decreased actin filament network formation, interrupted peristalsis, blocked ER expansion, and reduced ER-mitochondria association, while increasing reactive oxygen species generation and inducing premature epithelial cell mitosis. This apparent loss of gut homeostasis was recapitulated by trypanosome infection and, conversely, sHSP silencing in infected insects increased parasite numbers. Taken together, our data reveal a paramount role for sHSPs in gut cell homeostasis following blood meals and support the hypothesis that sHSP expression down regulation represents an adaptive manipulation of the insect host by the parasite.

**Significance Statement:** This study demonstrates that sHSPs serve as critical regulators of digestive physiology and intestinal homeostasis in a Chagas disease vector. These proteins orchestrate key processes including ROS production and cytoskeletal and endoplasmic reticulum organization, directly influencing *T. cruzi* proliferation. Conversely, the parasite manipulates their expression, revealing sHSPs as novel determinants of vector competence. Between 6 and 7 million people worldwide suffer from Chagas Disease, with 10,000 deaths annually. Although vector-borne transmission has declined, oral transmission is increasing, underscoring the urgent need to understand molecular factors governing vector competence. Our findings position sHSPs as promising targets for developing innovative strategies to control disease transmission.

## Introduction

Blood-sucking insects are capable of ingesting volumes of blood many times their own body weight, often in a very short period. This nutritional overload involves the intake of major components of vertebrate blood—such as water, salts, amino acids, iron, and heme—in amounts that would be toxic to most organisms (1). These threats are multifaceted: osmotic, as blood is nearly 80% water; oxidative, since iron and heme from hemoglobin degradation promote the formation of oxygen free radicals; metabolic, due to the large quantities of amino acids produced from protein digestion; and microbiological, as the gut microbiota can proliferate a hundred- to a thousand-fold within hours or days. Consequently, the adaptation to hematophagy is intrinsically linked to the evolution of robust stress response mechanisms that mitigate or prevent cellular damage (2, 3). Furthermore, pathogens ingested with the blood must navigate these host homeostatic responses to establish a successful infection and complete their life cycle.

Triatomine insects, or “kissing bugs,” are obligate hematophagous insects throughout their five nymphal instars, requiring at least one blood meal to trigger each molt (4). Several triatomine species are vectors of Chagas disease, an illness first described by Carlos Chagas in 1909, caused by the protozoan parasite *Trypanosoma cruzi* (5, 6). The *T. cruzi* life cycle within the insect vector is confined to the digestive tract. Following a blood meal, the number of circulating trypomastigotes is drastically reduced. However, after differentiating into the insect-specific epimastigote form – the replicative stage in the vector – the parasite population expands. A few days later, transformation into infective metacyclic trypomastigotes completes the parasite’s development in the insect, allowing for transmission to a new vertebrate host, either through contaminated feces or orally via ingestion of an infected insect (7). Although early reports considered *T. cruzi* subpathogenic to its vector (8, 9), a growing consensus now regards the parasite as mildly pathogenic, with clear signs of virulence manifesting primarily under stressful environmental conditions (10, 11). However, the underlying mechanisms of this pathogenicity remain largely uncharacterized. Moreover, molecular studies of triatomine immunity have shown that *T. cruzi* infection elicits an immune response and that host immune effectors control parasite populations, indicating that the parasite is recognized and managed as a pathogen by the insect’s immune system (12).

The cellular stress response is a conserved set of mechanisms that allows organisms to cope with deviations from homeostasis in a changing environment (13, 14). These pathways are also integral to normal physiological processes, such as development and cellular turnover (15). A hallmark of this response is the induction of protein chaperones, which counteract the protein misfolding and aggregation that are common consequences of cellular stress (13). Among these, the small Heat Shock Proteins (sHSPs) are a ubiquitous, yet often overlooked, multigenic family found in all three domains of life. sHSPs are ATP-independent molecular chaperones critical for maintaining protein homeostasis. Their chaperone activity stems from a unique, conserved alpha-crystallin domain of about 80 amino acids in the C-terminal region, which is flanked by highly variable sequences that facilitate dimerization, another conserved feature of these proteins (16, 17). Under stress, they bind to misfolded proteins, preventing their irreversible aggregation and facilitating their refolding by ATP-dependent chaperones (18). However, several studies have demonstrated that sHSPs are also active under non-stress conditions, where they maintain the integrity of cytoskeletal proteins (19, 20) and prevent the aggregation of actin and intermediate filaments during oxidative stress (21, 22). Thus, they function not only as stress-responsive agents but also as stress-preventive mechanisms.

Recently, a transcriptomic analysis of the digestive tract of first-instar *Rhodnius prolixus* nymphs revealed the involvement of several stress-response genes during both blood digestion and infection. Notably, this analysis highlighted members of the sHSP family that were modulated by both the blood meal and *T. cruzi* infection (23). Here, we investigate the role of these gut-expressed sHSPs in intestinal homeostasis following a blood meal and during *T. cruzi* infection, exposing their function as a target of the parasite’s virulence toward its insect host.

## Results

### Small heat-shock genes are modulated by blood meal and parasite infection

To identify genes modulated by blood feeding and *T. cruzi* infection, we analyzed a transcriptome generated from the midgut cDNA of first-instar nymphs under three conditions: unfed, fed with sterile blood, or fed with blood containing 10³ trypomastigotes/mL. This parasite concentration mimics the average parasitemia found in chronically infected vertebrate hosts. Furthermore, trypomastigotes are the parasitic form that the vector acquires during a natural blood meal. The analysis identified 8,046 differentially expressed genes (p < 0.05) in response to either blood feeding or parasite infection (23).

Among these, five sHSP genes were expressed in the gut, all of which were differentially regulated by feeding and infection. In blood-fed nymphs, two genes (RP004515 and RP008537) were upregulated throughout the post-meal period, while the other three (RP005869, RP011567, and RP015293) were upregulated only on the first day after feeding (Figure 1A). Remarkably, *T. cruzi* infection downregulated all five of these sHSP genes. In addition to these five, a search for proteins containing the α-crystallin domain (PF00011) identified three other sHSP paralogs in the *R. prolixus* genome (RPRC005139, RPRC009973, and RPRC013971) that were not expressed in the gut (Figure 1B). Phylogenetic analysis revealed that four of the five gut-expressed transcripts clustered together with 65-85% identity, while RPRC008537 showed only 35% identity. The three non-digestive paralogs exhibited less than 25% identity, suggesting a recent gene expansion event produced the forms expressed in the gut (Figure 1C).

**Figure 1.**
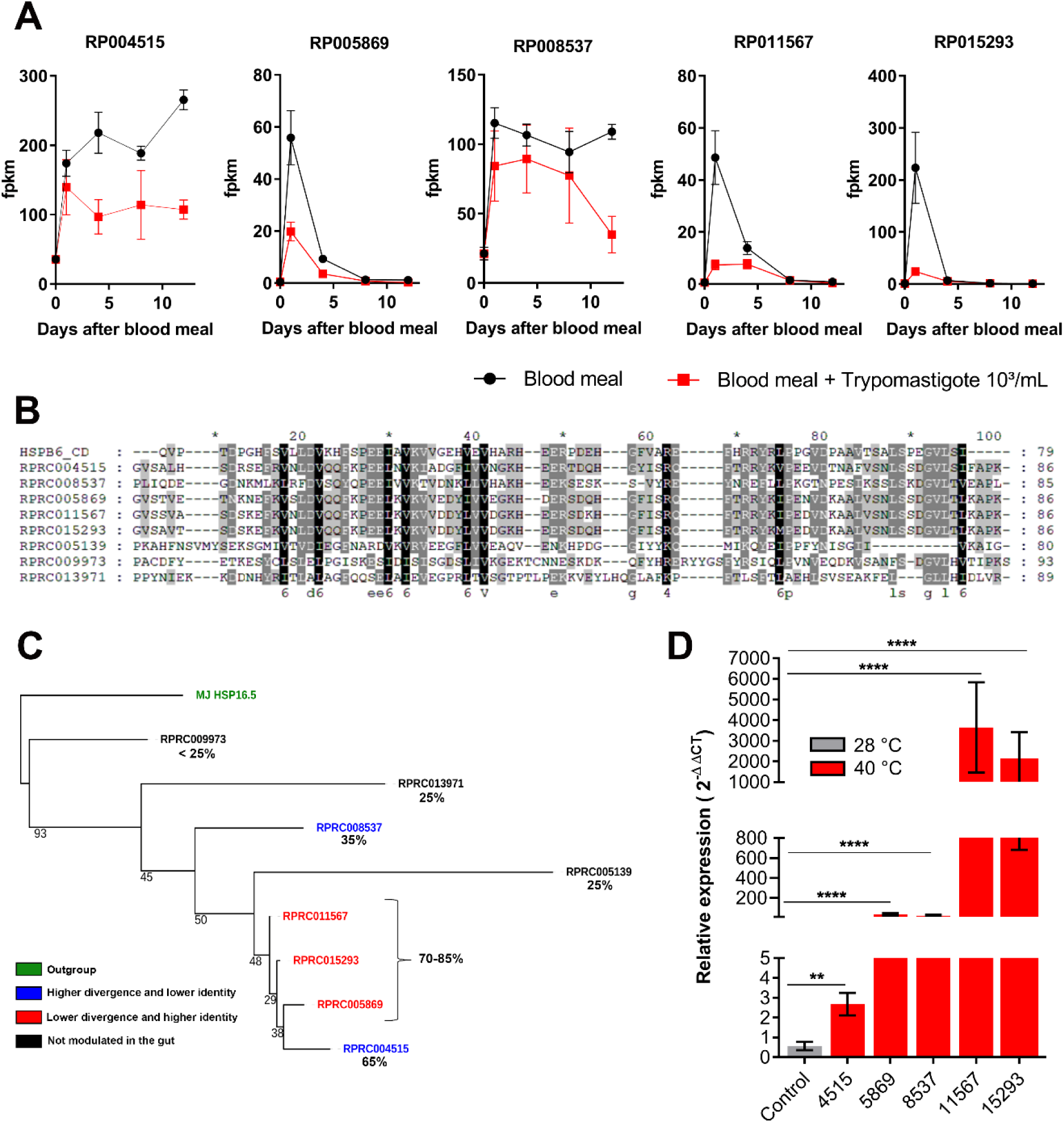
Small Heat-Shock (sHSPs) genes showed increased expression in the insect gut after a blood meal and are inhibited by parasite infection. (A) Five sHSPs were upregulated after a blood meal (black circles) and were downregulated by an infected blood meal (*T. cruzi* CL-Brenner clone, 10^3^ trypomastigote parasites/mL of blood; red squares). Shown are data from a transcriptome of *Rhodnius prolixus* midgut (23). (B) Alignment of sHSPs sequences found in the kissing bug genome with a consensus sequence of the conserved α-crystalline domain. Alignments were performed in Seaview software using MUSCLE sequence alignment program. (C) Phylogenetic analysis of *R. prolixus* sHSPs sequences was performed using the conserved α-crystalline domain (PFAM - PF00011) PhyML, with the JTT substitution model and 500 bootstrap replicates. A sHSP sequence of a thermophilic archaeon *Methanocaldococcus jannaschii* (Hsp 16.5 – NCBI Accession number : WP_010869783.1) was used as an outgroup. (D) sHSPs response to heat shock - Four days after feeding, the first-instar nymphs were exposed to 40°C for 2 hours, followed by 2 hours back at 28°C. Expression of the sHSPs was determined by qPCR, relative to controls kept at 28°C. Data shown are mean ± SEM, (n=3 and individual samples were pool of 10 insects). Statistical analysis with T-test calculated with the control of each group.

This expression pattern was confirmed by qPCR performed under the same experimental design as the transcriptome analysis. The only exception was gene RP11567, which did not show the pronounced decrease in expression on the first day post-infection that was observed in the transcriptomic data (Figure S1).

The response to a heat shock is the archetypal feature that gives the sHSP protein family its name, leading us to test if these genes were responsive to thermal stress. Measurement of the expression of the five gut-expressed sHSPs in first-instar nymphs following a heat shock showed that four of the sHSP transcripts (RP008537, RP005869, RP011567, and RP015293) were strongly upregulated, with expression increasing by 20- to 4,000-fold compared to control insects (Figure 1D).

### sHSPs play a paramount role in intestinal cell physiology and homeostasis by controlling redox status and actin filament assembly

Given that all five sHSPs expressed in the midgut are induced by a blood meal and downregulated by infection, we chose to collectively knock down these genes to elucidate their role in gut physiology. A previous study showed that injecting dsRNA into adult females results in offspring with persistent and efficient gene silencing (24). Therefore, we injected a mixture of dsRNAs targeting the five sHSPs (hereafter referred to as dsMix) into adult females, using the parental RNAi approach to generate first-instar nymphs with silencing efficiencies ranging from 50% to 99% (SI Appendix, Figure S2). Importantly, several key aspects of insect biology remained unaffected: there was no reduction in adult female oviposition, egg hatching, or the rate of blood protein digestion in silenced first-instar nymphs (SI Appendix, Figures S3 and S4).

sHSPs function as stress response proteins not only through their chaperone activity (17) but also through their roles in oxidative stress resistance and cell death prevention (25–29). Silenced nymphs exhibited a marked increase in DHE fluorescence – a ROS-sensitive probe - after a blood meal in the posterior midgut, indicating elevated ROS generation. Importantly, this fluorescence increase was prevented by incubation with MitoTEMPO, a selective mitochondrial antioxidant, implicating gut epithelial mitochondria as a major source of the ROS observed upon sHSP silencing (Figures 2A and 2B).

**Figure 2.**
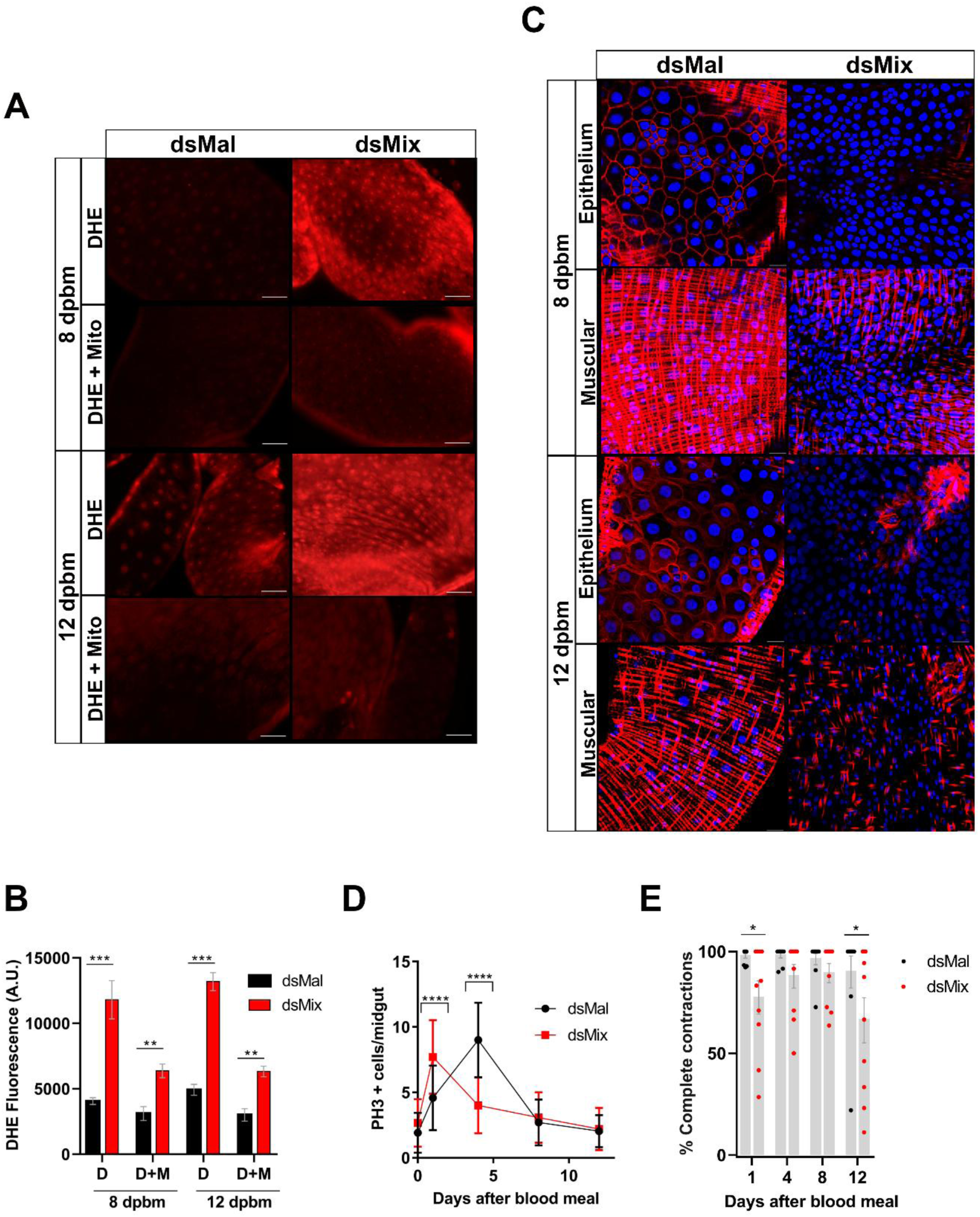
sHSPs knockdown affects ROS production, actin network, intestinal homeostasis and contractility. A) ROS levels were evaluated by DHE oxidation in the absence or presence of MitoTEMPO 5 µM (DHE+Mito), an antioxidant selective for mitochondrial ROS. Shown are representative images of posterior midgut of first-instar nymphs taken at 8 and 12 days post-blood meal (scale bar: 10µM). (B) Quantitative analysis of ROS levels from fluorescence DHE oxidation signal in the absence (D) or presence of MitoTEMPO (D+M), using ZEISS Axio Observer software (data shown are mean ± SEM; n = 3). (C) Phalloidin (red) and DAPI (blue) staining of the anterior midgut of control (dsMal) or sHSPs-silenced first-instar nymphs (dsMix). (D) Epithelial cell proliferation was assessed as the number of phosphorylated histone H3-positive (PH3+) cells in the midgut of control (dsMal) and sHSPs-silenced (dsMix) insects (data shown are mean ± SEM; n= 3). (E) Peristaltic contraction profile of the anterior midgut of nymphs silenced for sHSPs was visually analyzed and the frequency of interrupted peristaltic waves was evaluated. Data shown are mean ± SEM (n = 10). Asterisks indicate p< 0.05 using a t-test calculated against each group’s respective control.

ROS-induced oxidative stress is a well-known inducer of cell death (30), and digestive epithelial cells exhibit the highest cellular turnover rates among all metazoan tissues. Therefore, an immediate regenerative response based on cell proliferation is the hallmark of gut homeostasis (31). Cell proliferation has been associated with responses to stress and tissue damage in both *Drosophila* and mice (32). Quantification of proliferative cells (phospho-histone H3 positive) revealed that while an increase in mitotic epithelial cells normally occurs by day 4 after feeding, this peak was anticipated upon sHSP silencing (Figure 2D).

In *Drosophila*, sHSPs are important not only for proteostasis control but also for cytoskeletal regulation through binding to actin, leading to stabilization of the actin filament network (20, 26). Sequence alignment of *R. prolixus* sHSPs revealed the presence of actin-binding domains similar to those found in filamin and alpha-actinin, which could facilitate F-actin filament cross-linking (33) (SI Appendix, Figure S5). Phalloidin staining of the midgut demonstrated a profound impact of sHSP knockdown on actin filament network formation in both epithelial cells and the surrounding muscular layer, leading to near-complete cytoskeletal disruption in both tissues (Figure 2C). After a blood meal, the anterior midgut showed strong periodic waves of contraction that moved anteroposteriorly along the entire organ, from the esophagus to the junction with the posterior midgut. Consistent with the effects on actin network of the muscular layer, sHSP silencing altered the contraction pattern of the anterior midgut, leading to interrupted wave propagation, frequently at the median portion of the gut, revealing that sHSPs are essential for normal contractile activity (Figure 2E and Supplementary movie files S1-4).

These results suggested a loss of intestinal homeostasis upon sHSP silencing, prompting us to analyze the ultrastructure of posterior midgut epithelial cells. In control insects (dsMal – Figure 3A), the apical region near the microvilli contained abundant mitochondria (Figure 3C) that were frequently associated with dilated endoplasmic reticulum (ER) cisternae (Figure 3B). In contrast, sHSP-silenced insects (Figure 3D) displayed the classical stacked sheet morphology of ER cisternae and lacked the intimate mitochondrial association observed in control cells (Figures 3E and 3F). ER expansion forming dilated cisternae is typically associated with cellular stress responses, yet here it was unexpectedly observed as part of normal physiology. Since these ER morphological changes usually accompany cell-protective mechanisms, we hypothesized that ER expansion observed in control insects represents a stress-preventive adaptation (see Discussion).

**Figure 3.**
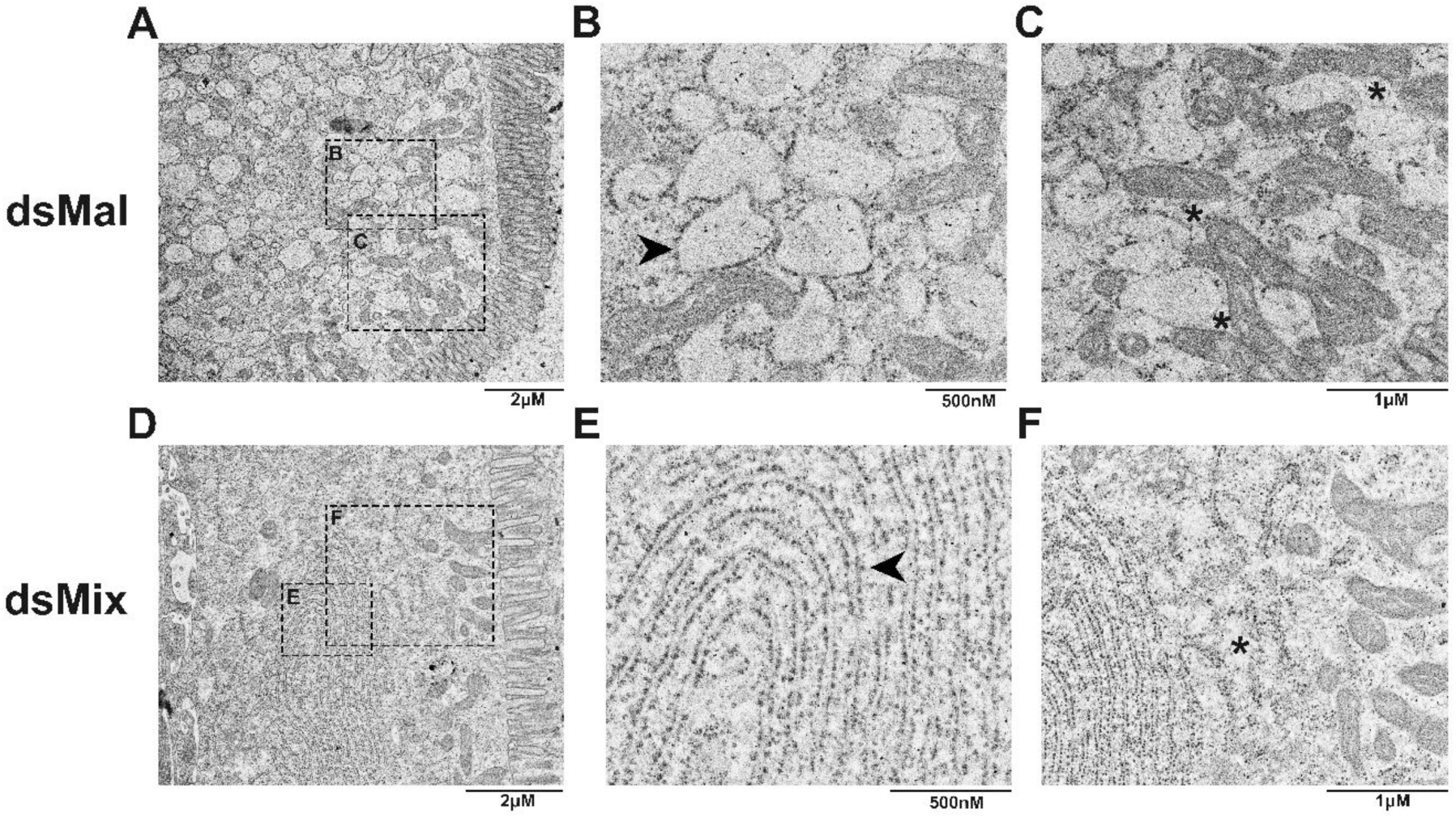
sHSP silencing changes endoplasmic reticulum (ER) organization. Show are transmission electron microscopy images of posterior midgut epithelial cells from control first-instar nymphs (A-C; dsMal) compared with sHSP-silenced nymphs (D-E; dsMix) revealing marked morphological differences in the ER (indicated by black arrows in B; dsMal and E; dsMix). Control insects display extensive dilation of ER cisternae, along with a close association between dilated ER and mitochondria (black asterisks in C; dsMal), contrasting with the lack of interaction found in silenced insects (black asterisk in F; dsMix).

### Multiple sHSP silencing increases *T. cruzi* burden and impacts intestinal physiology

Since our transcriptomic analysis showed that parasite infection downregulated sHSP expression, we tested whether sHSP silencing would affect parasite numbers. Quantitative evaluation of *T. cruzi* by qPCR demonstrated that sHSP silencing (dsMix) increased parasite populations following an infected blood meal throughout the entire study period (Figure 4A).

**Figure 4.**
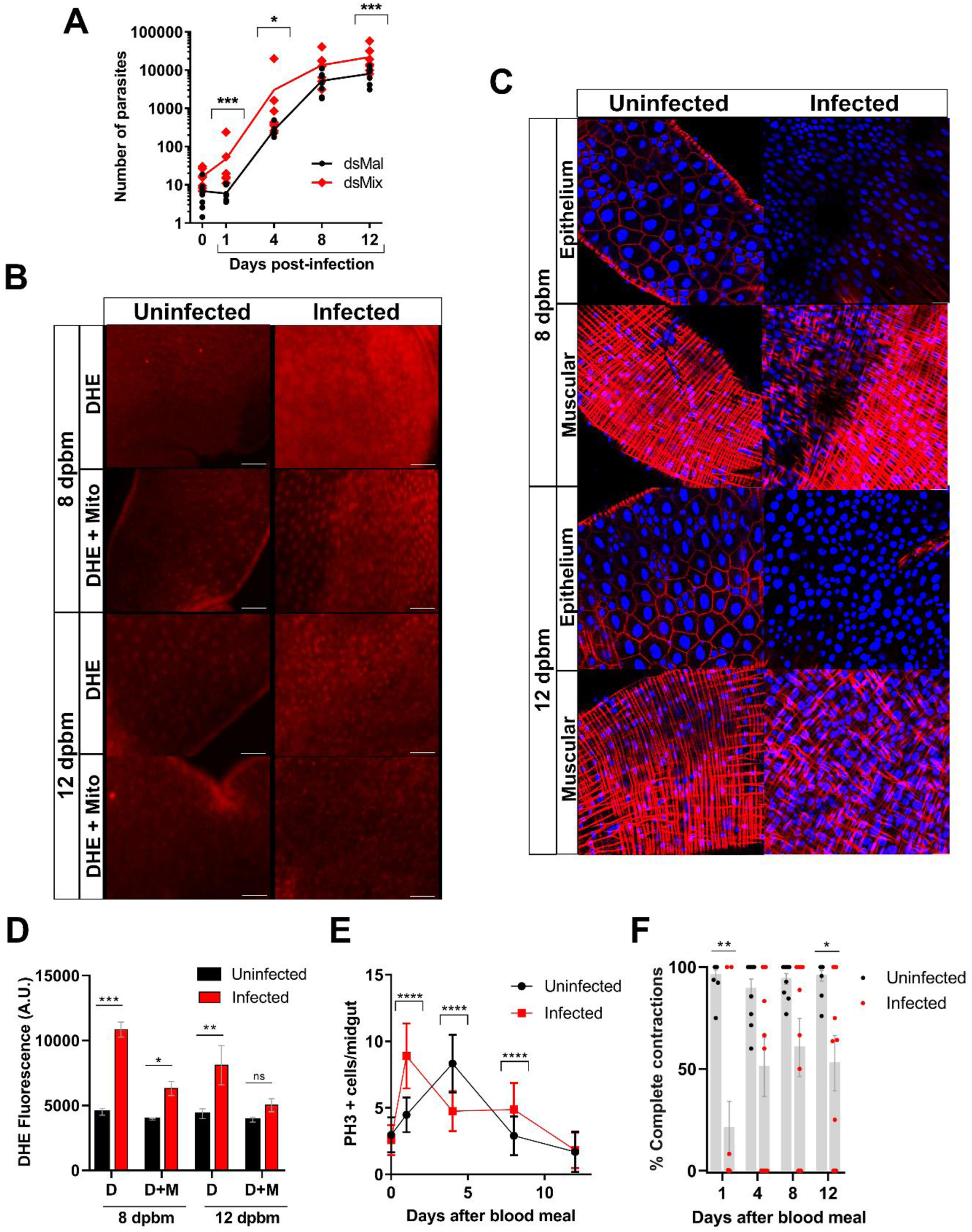
sHSP knockdown increases *Trypanosoma cruzi* proliferation in the insect gut and infection recapitulates sHSP silencing phenotype. (A) Parasite number after an infective blood meal. (B) DHE staining of posterior midgut to evaluate ROS levels upon infection, in the absence or presence of 5 µM MitoTEMPO. Shown are representative images of fluorescence micrographs taken at 8 and 12 days post-blood meal (scale bar: 10µM). C) Phalloidin (red) and DAPI (blue) staining of the anterior midgut of first-instar nymphs at the epithelial and muscular layers. (D) Quantitative analysis of ROS levels from fluorescence DHE oxidation in the absence (D) or presence of MitoTEMPO (D+M). Data shown are mean ± SEM; n= 3). (E) Cellular proliferation assessed as of the number of phosphorylated histone H3-positive (PH3+) cells. Data shown are mean ± SEM; n= 3. (F) Peristaltic contraction profile of the anterior midgut of control (black dots) and infected insects (red dots) showing the frequency of interrupted peristaltic waves. Images were taken at several days after a blood meal Data shown are mean ± SEM (n = 10). Asterisks indicate statistical difference, t-tests relative to each group respective control (* p<0.05; ** p<0.01; *** p<0.001).

The impact of RNAi-mediated sHSP reduction on intestinal homeostasis led us to hypothesize that sHSP downregulation by *T. cruzi* infection would similarly disrupt host gut physiology while promoting parasite proliferation. Measurement of ROS levels, mitotic activity, and actin filament networks in insects receiving an infected blood meal recapitulated the phenotypes observed upon sHSP gene silencing, including increased ROS production in the posterior midgut (prevented by MitoTEMPO), enhanced epithelial cell proliferation, and disrupted actin network organization in both muscular and epithelial layers and altered contractile activity (Figures 4B-F). For most parameters observed, phenotypes produced by infection alone (Figure 4) were relatively milder compared to those resulting from infection combined with sHSPs (dsMix) gene silencing (Figure 2).

## Discussion

Cellular responses to stressful conditions are fundamental to the capacity of living cells to cope with an ever-changing environment. Members of the sHSP family are ubiquitous chaperones that are constitutively expressed across different cell types and tissues (17) and participate in both stress prevention and response mechanisms (25). Blood-sucking insects face an enormous physiological challenge by taking blood meals several times their body weight, and the capacity to maintain homeostasis despite this exceptionally large nutrient influx has been proposed as a major adaptation to hematophagy. Here, we characterized the *Rhodnius prolixus* multigene sHSP family, revealing their essential role in maintaining intestinal homeostasis following blood meals. Mechanistically, our data indicate that sHSPs provide at least four key functions in digestive apparatus physiology: they are critical for actin network assembly in both epithelial and muscular layers, for epithelial cell renewal, and for coping with cellular stressors that come along with the blood meals. We also demonstrate that the human pathogen *Trypanosoma cruzi* manipulates host sHSP expression for its own benefit, enhancing its proliferation at the cost of disrupting intestinal homeostasis through actin network compromise and oxidative stress induction.

The effects of sHSP gene silencing on actin filament formation, which are recapitulated by *T. cruzi* infection, likely affect multiple aspects of digestive physiology. Notably, sHSP silencing compromised the contractile capacity of midgut-enclosing muscular tissues and impaired peristalsis (Figures 2E and S4). Tissues involved in mechanical work are inherently subjected to mechanical stress, as movements impose direct physical damage that can be attenuated and repaired through chaperone action. Several reports document sHSP interactions with mechanosensitive proteins that regulate contractile activity (19, 26). Although not experimentally addressed here, epithelial cytoskeletal disorganization likely also affects the mechanical stiffness of the epithelium that underlies gut wall barrier function (34).

Transmission electron microscopy of normal insects revealed dilated ER cisternae in epithelial cells (Figure 2E-G), an unexpected finding since this ER morphology typically occurs in stress-exposed cells (35–37). These observations align with reports by Billingsley and Downe from the 1980s, documenting the presence of apical mitochondrial clustering and dilated ER after feeding in *Rhodnius* midgut epithelial cells (38, 39). As we have noted, blood meals in hematophagous insects expose gut tissues to multiple potential stressors, including overloads of heme and iron – pro-oxidant molecules produced from dietary hemoglobin hydrolysis (2, 3). Moreover, these insects consume blood volumes several times their body weight, and vertebrate blood is 87% protein on a dry-weight basis, requiring enormous metabolic effort for synthesizing and exporting digestive proteins. This represents a potential ER stress source, typically associated with oxidative stress and unfolded protein response (UPR) activation. Interestingly, ER expansion has been shown to alleviate ER stress, particularly in specialized secretory cells handling large cargo loads, as expanded ER accommodates increased client proteins, reduces folding intermediate concentrations, and decreases protein aggregation likelihood (40). Therefore, the ER expansion forming dilated cisternae shown in Figure 3A may represent epithelial cells activating a stress-preventive response during digestion. Conversely, the canonical stacked-sheet ER cisternae positioned away from mitochondria in sHSP-silenced insects (Figure 3B) indicates failure to activate this preventive ER stress program.

Another feature that is frequently observed together with ER expansion is the close association between dilated ER cisternae and mitochondria (41). Both mitochondria and ER are major sites of reactive oxygen species (ROS) production, and mitochondria-ER contact sites play pivotal roles in regulating eukaryotic cell redox balance (41, 42). ER expansion occurs under both stress conditions (43–45) and homeostatic conditions, such as during mitosis or intense secretory activity, where close apposition between enlarged ER cisternae and mitochondria has been reported (41). Importantly, we demonstrated that sHSP silencing prevented mitochondria-ER association, markedly reducing contact sites between these organelles (Figure 3) while simultaneously increasing mitochondrial ROS production. While these findings align with previous literature observations in different models, our results implicate sHSPs as important upstream players in pathways leading to ER expansion and mitochondria-ER colocalization.

Regarding the increased ROS levels observed here, data from both human and *Drosophila* cells demonstrate that HSP27 – an sHSP family member – promotes ROS resistance by increasing glutathione levels (29). Interestingly, recent data show that sHSP HSPB1 is a chaperone found also in the mitochondrial intermembrane space (46). Notably, HSP27 also modulates actin filament integrity, contributing to cytochrome c retention in mitochondria, thus linking cytoskeletal damage to mitochondrial dysfunction and apoptosis (47). Therefore, the prevention of mitochondria-derived oxidative stress observed here could result from direct sHSP participation in this organelle. Alternatively, actin filament disorganization induced by sHSP silencing could indirectly prevent ER expansion and hinder the ER-mitochondria coupling needed to limit ROS levels.

This scenario fits our data showing *T. cruzi*-elicited sHSP repression (emulated by RNAi gene silencing) causing a syndrome encompassing actin filament disorganization, increased ROS, and accelerated regenerative responses based on intestinal stem cell recruitment. The only previous work on heat shock proteins in triatomine physiology demonstrated that HSP70 chaperone knockdown in the gut affected blood processing, causing digestive delays, energy metabolism impairment, and immune gene modulation (48). In contrast, we observed that sHSP knockdown does not affect insect survival or digestion.

sHSPs are scarcely mentioned in insect disease vector literature. Notably, sHSPs were among the most highly upregulated transcripts in *Aedes aegypti* cells exposed to heme (49). Interestingly, a search for commonly regulated genes in *Aedes aegypti* responses to four different arbovirus infections identified five sHSPs among the 11 genes upregulated in all cases (50). Recently, Qu et al. (2023) (51), working with cell culture lines, suggested that Chikungunya infection upregulates sHSP expression and observed increased viral infection upon sHSP silencing. Our data suggest that sHSP downregulation by *T. cruzi* represents host manipulation that enhances parasite fitness by promoting multiplication in the intestinal extracellular space. Interestingly, elevated ROS production is almost invariably described as a pathogen-killing mechanism. However, *T. cruzi* may be among the few exceptions, as it has been demonstrated that ROS production benefits the parasite, with oxidative environments contributing to both trypanosome persistence in vertebrate hosts (52) and improved epimastigote – the insect developmental stage – proliferation (53). *R. prolixus* sHSP expression is promptly triggered by blood meals, though the regulatory mechanism remains unknown. Phylogenetic analysis indicates recent gene expansion (Figure 1C) producing gut-expressed sHSPs, which could speculatively indicate recruitment of a specific paralog functioning as a homeostatic hub facilitating insect gut responses to incoming blood meals. Taken together, our results suggest a role for sHSPs in adaptation to hematophagy, revealing essential participation of this protein family in maintaining intestinal homeostasis. Moreover, this protein family emerges as a molecular target for parasite manipulation, constituting a pathogenicity factor toward the insect vector digestive apparatus.

## Materials and Methods

### Insects

*Rhodnius prolixus* were maintained at 28°C with 70-80% relative humidity under a 12-hour light/dark photoperiod. Insects were fed on rabbit blood. All animal handling procedures followed protocols approved by the Ethics Committee on Animal Use at UFRJ (registration #123/22). Insects used here were first-instar nymphs (N1).

### Parasites

*Trypanosoma cruzi* clone CL-Brener was propagated in LLC-MK2 cells maintained in DMEM supplemented with 10% Fetal Bovine Serum (FBS) and antibiotics (Penicillin 100 U/mL + Streptomycin 100 μg/mL). For infections, cells were transferred to fresh bottles containing DMEM with 2% FBS and antibiotics, and parasites were added at approximately 1.65 parasites per cell. After 48 hours, culture supernatant was centrifuged at 1,500 rpm for 5 minutes, the pellet discarded, and the supernatant re-centrifuged at 3,000 rpm for 15 minutes. The parasite-containing pellet was resuspended in 1 mL DMEM + 2% FBS, counted using a Neubauer chamber, and maintained at 37°C for up to 3 hours before use.

### *In silico* analysis

sHSP sequences in the *Rhodnius prolixus* genome were identified using FAT (Functional Analysis Tool) software (MESQUITA et al., 2011) by searching for the alpha-crystallin conserved domain (PF00011 – PFAM). The five sHSP sequences analyzed (RPRC004515, RPRC008537, RPRC005869, RPRC011567, and RPRC015293) were identified from our transcriptomic analysis (Contreras et al., 2015; unpublished) as transcripts modulated by blood feeding and parasite infection in first instar nymphs, and were mapped to the *Rhodnius prolixus* genome in the VectorBase database (Genome version: RproC3.3 - https://www.vectorbase.org/).

Sequence alignments were performed using SEAVIEW software (54) with the MUSCLE multiple sequence alignment method (55). Phylogenetic analysis utilized only the conserved alpha-crystallin domain regions. Trees were generated in SEAVIEW using PhyML with the JTT model and default parameters, with bootstrap analysis (500 replicates) used to assess branching support.

### RNA extraction, cDNA synthesis, and quantitative RT-PCR

For gene silencing experiments and sHSP expression analysis, dissected intestines were transferred to microtubes containing 300 μL RNA extraction solution [38% saturated phenol (pH 4.3), 800 mM guanidine thiocyanate, 400 mM ammonium thiocyanate, 100 mM sodium acetate (pH 5.0), 5% glycerol] (56). Intestines were macerated in extraction solution, then 100 μL of chloroform was added and samples were vortexed for 20 seconds. Following centrifugation at 12,000×g for 10 minutes at 4°C, the aqueous phase was transferred to a fresh tube and mixed 1:1 (v/v) with isopropanol. After vortexing for 20 seconds and 10-minute room temperature incubation, samples were centrifuged at 14,000 rpm for 10 minutes at 4°C to pellet RNA. The supernatant was removed and RNA pellets were washed with 1 mL 70% ethanol, re-centrifuged at 14,000 rpm for 10 minutes at 4°C, then washed again with 100% ethanol. After ethanol removal and drying, RNA was resuspended in 10 μL sterile DEPC-treated water and quantified using a Nanodrop spectrophotometer (ThermoScientific). For cDNA synthesis, 1 μg RNA was treated with DNase for 30 minutes at 37°C, then the enzyme was inactivated at 65°C for 10 minutes with 1.25 mM EDTA. cDNA synthesis was performed using the High-Capacity cDNA Reverse Transcription kit (Applied Biosystems) following manufacturer’s instructions.

Gene expression analysis by qPCR employed a reaction mix containing 2× PCR buffer without MgCl₂ (Sigma), 5 nM MgCl₂ (Sigma), 2× SYBR Green I, 1× quantitative PCR reference dye (Sigma), 400 nM dNTP mix, and 0.5 U/μL G2 Go Taq Polymerase Hot Start™ (ProMega) (57). Reactions contained 7.5 μL reaction mix, primer mix, DEPC-treated ultrapure water to 10 μL, and 5 μL cDNA (1:10 dilution) for a final volume of 15 μL. Oligonucleotide sequences and concentrations are listed in Table 1 (Supporting Information). Reactions were performed using StepOnePlus Real Time PCR equipment (Applied Biosystems) and analyzed with StepOne software. Relative gene expression was calculated using the ΔCt method (58) to analyze transcriptome validation data and ΔΔCt method (58) to analize knockdown and heat shock data, with EF-1 (Elongation factor-1) as the endogenous control gene (59).

### dsRNA synthesis and parental gene silencing

Double-stranded RNA synthesis utilized T7 RNA polymerase expressed from plasmid Pt7-911 (provided by Dr. Thomas E. Shrader, Albert Einstein College of Medicine) (60). dsRNA transcription was adapted from an established tRNA transcription (61). Briefly, reactions were performed at 37°C for 16 hours in medium containing 40 mM Tris·HCl (pH 8.0), 22 mM MgCl₂, 5 mM DTT, 2 mM spermidine, 0.05% BSA, 15 mM guanosine monophosphate, 7.5 mM each nucleoside triphosphate, PCR-amplified cDNA from fed first-instar nymph whole bodies (0.1 μg/μL), and 5 μM T7 RNA polymerase. dsRNA transcripts were treated with 1 μL DNase (2 U/μL) at 37°C for 30 minutes, then precipitated using 1:10 (v/v) 3 M sodium acetate (pH 5.2) and 1:1 (v/v) isopropanol. Pellets were washed twice with 70% ethanol, resuspended in water to 1 μg/μL, and quantified using a Nanodrop 1000 v.3.7 spectrophotometer (ThermoFisher Scientific). *Escherichia coli* maltose binding protein gene (dsMal, Gene ID: 948538) served as the nonspecific control dsRNA. Oligonucleotide sequences and concentrations are listed in Table S2. Adult females in their second or third feeding/oviposition cycle were used for silencing experiments. dsRNA injection occurred on day 21 post-feeding using a 10 μL Hamilton syringe to deliver dsRNA directly into the hemocoel via the ventral region at the third thoracic-abdominal segment junction. Each female received 1 μL dsMal or experimental dsRNA (both at 1 μg/μL). Six days post-injection, females were fed on rabbit ears and individually housed in glass flasks with filter paper to monitor oviposition and hatching. Five to seven days after hatching, nymphs were fed with blood, and after 4 days, intestinal dissection, RNA extraction, and parental silencing analysis by qPCR were performed.

### Heat shock treatment

Four days post-feeding, first-instar nymphs were subjected to heat shock at 40°C for 2 hours, then returned to 28°C for 2 hours recovery (48) before digestive apparatus dissection and sHSP expression evaluation by qPCR. Control insects remained at 28°C throughout.

### Blood digestion time course

Blood meal digestion kinetics were assessed by measuring total protein content of digestive apparatus from insects dissected at various post-feeding timepoints. Protein quantification was performed using the Lowry method (62) with bovine serum albumin as standard using a plate spectrophotometer (SpectraMax M3 – Molecular Devices).

### Reactive Oxygen Species (ROS) detection

For evaluation of ROS levels, midguts of first-instar nymphs were dissected and incubated in Leibovitz-15 media supplemented with 5% FBS (L15/5%), then transferred to L15/5% containing 25 μM dihydroethidium (hydroethidine; DHE; Invitrogen), an oxidant-sensitive fluorophore. Samples were incubated in the dark at 28°C for 25 minutes, washed with Phosphate Buffered Saline (0.15 M NaCl, 10 mM sodium-phosphate buffer, pH 7.2; PBS), and immediately mounted on glass slides for fluorescence microscopy. In the MitoTEMPO experiments, after incubation with DHE, the samples were additionally incubated with 5 µM MitoTEMPO ((2-(2,2,6,6-Tetramethylpiperidin-1-oxyl-4-ylamino)-2-oxoethyl) triphenylphosphonium chloride; Sigma Aldrich) in the dark at 28 °C for 25 minutes, followed by washing steps performed as described above. Images were acquired using a Zeiss Observer.Z1 equipped with Zeiss Axio Cam MrM and a filter set used for DHE labeling (excitation BP 546/12 nm; beam splitter FT 580 nm; emission LP 590 nm) with 20× objective and 80 ms exposure. For each insect, 20 different fields were acquired along the length of the midgut segment, and the average intensity value was used for quantitative analysis. Quantitative fluorescence analysis employed AxioVision version 4.8 software.

### Actin filament staining

For actin visualization, first-instar nymph midguts were dissected in PBS and fixed in 4% paraformaldehyde for 30 minutes. Samples were washed twice with PBS (10 minutes each) and permeabilized with 0.1% Triton X-100 in PBS for 15 minutes. Following permeabilization, samples were incubated with 200 μL Alexa Fluor™ 546 Phalloidin (ThermoFisher Scientific) diluted 1:1000 (v/v) in PBS for 40 minutes at 28°C. After two PBS washes, samples were mounted using VECTASHIELD® Antifade Mounting Medium with DAPI (Vector Laboratories). Representative images were acquired using a Leica SP5 confocal laser-scanning inverted microscope with 40× objective and processed using LAS X software.

### Cell proliferation analysis

For identification and quantification of mitotic cells, midguts were dissected in PBS, followed by fixation in paraformaldehyde 4% for 30 minutes. Samples were washed twice with PBS (10 minutes each) and permeabilized with 0.1% Triton X-100 in PBS for 15 minutes. After 30-minute blocking in solution containing 0.1% Tween 20 and 5% BSA in PBS at room temperature, samples were incubated overnight with primary mouse anti-PH3 antibody (Merck Millipore) at 1:1000 (v/v) dilution in blocking solution. Following three 20-minute washes in 0.1% Tween 20, 0.25% BSA in PBS, samples were incubated with secondary goat anti-mouse antibody conjugated to Alexa Fluor 546 (ThermoFisher Scientific) at 1:2000 (v/v) dilution in blocking solution for 2 hours at room temperature. Final washes in 0.1% Tween 20, 0.25% BSA in PBS preceded mounting with VECTASHIELD® Antifade Mounting Medium with DAPI (Vector Laboratories). Images were acquired using Zeiss Observer Z1 with Zeiss Axio Cam MrM and analyzed using AxioVision version 4.8 software (Carl Zeiss AG).

### Experimental infection and *T. cruzi* quantification

Heparinized rabbit blood (0.5 U/mL) was centrifuged at 6,000 rpm, serum heat-inactivated at 56°C for 45 minutes, and erythrocytes washed three times in 0.15 M NaCl before reconstitution with inactivated serum. First-instar nymphs were artificially fed reconstituted blood (2.5 U/mL) containing 10³ trypomastigotes/mL from CL Brener strain. Post-feeding, insects were maintained at 28°C and samples were frozen at -80°C after dissection at specified timepoints. Only fully engorged animals were analyzed (61).

Parasite quantification was performed as described elsewhere (63). Insects were homogenized using plastic tissue grinders in 1 mL solution containing 1.5% CTAB, 2 M NaCl, 10 mM EDTA, 100 mM sodium acetate (pH 4.6), and 10 μg/mL salmon sperm DNA (Sigma-Aldrich). Plasmid Litmus 28i-mal (New England Biolabs) at 10 ng/mL served as internal heterologous reference for DNA recovery efficiency quantification. Extracted DNA was resuspended in 50 μL sterile deionized water.

Parasite quantification employed qPCR assays in 15 μL final volumes containing 7.5 μL qPCR mix (as described above), 5 μL extracted DNA, and 500 nM specific oligonucleotides targeting repetitive sequences in *T. cruzi* genomic DNA (64, 65). Parallel reactions used 5 μL plasmid Litmus 28i-mal DNA and 500 nM Mal gene-specific oligonucleotides as curve controls. Parasite numbers were calculated using standard *T. cruzi* DNA amplification curves and normalized against plasmid Litmus 28i-mal internal standard results. The oligonucleotides for qPCR used in this study are listed in Table SI 1.

### Peristaltic contractions

N1 nymphs were observed under a stereomicroscope, and contractions were recorded for 5 minutes using a cell phone camera (iPhone 13) attached to the stereomicroscope using a Celestron-NexYZ 3-Axis Smartphone Adapter. For video image acquisition, the nymphs were immersed in drop of water on a glass plate with the dorsal side upward and a coverslip was placed over the insects to keep them stable during recording, using a 2 mm O-ring as a spacer. Illumination was provided by LED light sources positioned laterally and above the setup. For each nymph, peristaltic contractions occurring within the 5-minute recording period were manually counted. A contraction was classified as complete when the peristaltic wave originated at the anterior end of the anterior midgut and propagated continuously to the posterior end.

### Transmission electron microscopy

Posterior midgut tissues were dissected 8 days after blood meal and immediately fixed by immersion in 2.5% glutaraldehyde (Grade I) and 4% freshly prepared formaldehyde in 0.1 M cacodylate buffer (pH 7.3). Samples were rinsed in cacodylate buffer, dehydrated through a graded acetone series, and embedded in Polybed 812 resin (Spurr). Ultrathin sections were stained with uranyl acetate and lead citrate, and subsequently examined using a Hitachi HT7800 TEM (operating at 100 kV). For each experimental condition, images from two individual insects were analyzed and at least ten images were acquired from each area of the tissue.

### Statistical Analysis

All experiments were performed in at least three biological replicates. The graphs and statistical analyses were generated using GraphPad Prism 8 software.

## Acknowledgments

We thank all members of the Laboratório de Bioquímica de Artrópodes Hematófagos for their critical suggestions, Leila Faustino for technical assistance regarding parasite culture, and Charlion Cosme and, S. R. Cássia for providing technical assistance and CENABIO-UFRJ for providing microscopy equipment and facilities. This work was supported by grants from FAPERJ, CAPES,and CNPq.

## Author Contributions

T.C.G.S, I.R, G.O.P.S, and P.L.O Conceived and designed the experiments. T.C.G.S, A.B.W.N, J.C.T.P and M.R.F Performed the experiments. T.C.G.S, A.B.W.N, J.C.T.P, M.R.F, F.A.D, H.D.P.C, I.R, G.O.P.S and R.D.M analyzed the data. H.D.P.C, F.A.D, I.R, G.O.P.S and R.D.M Contributed with previous data and analysis tools. T.C.G.S and P.L.O wrote the paper and conducted statistical analysis. All authors have read, revised and agreed to the published version of the manuscript.

