## Supplemental Information figures and tables for "Small Heat Shock Proteins have a Paramount Role in *Trypanosoma cruzi* Infection Impacting Intestinal Homeostasis of an insect vector of Chagas Disease"

**This PDF file includes:**

Supporting text
Figures S1 to S5
Tables S1 to S2
Legends for Movies S1 to S4
SI References

**Other supporting materials for this manuscript include the following:**

Movies S1 to S4

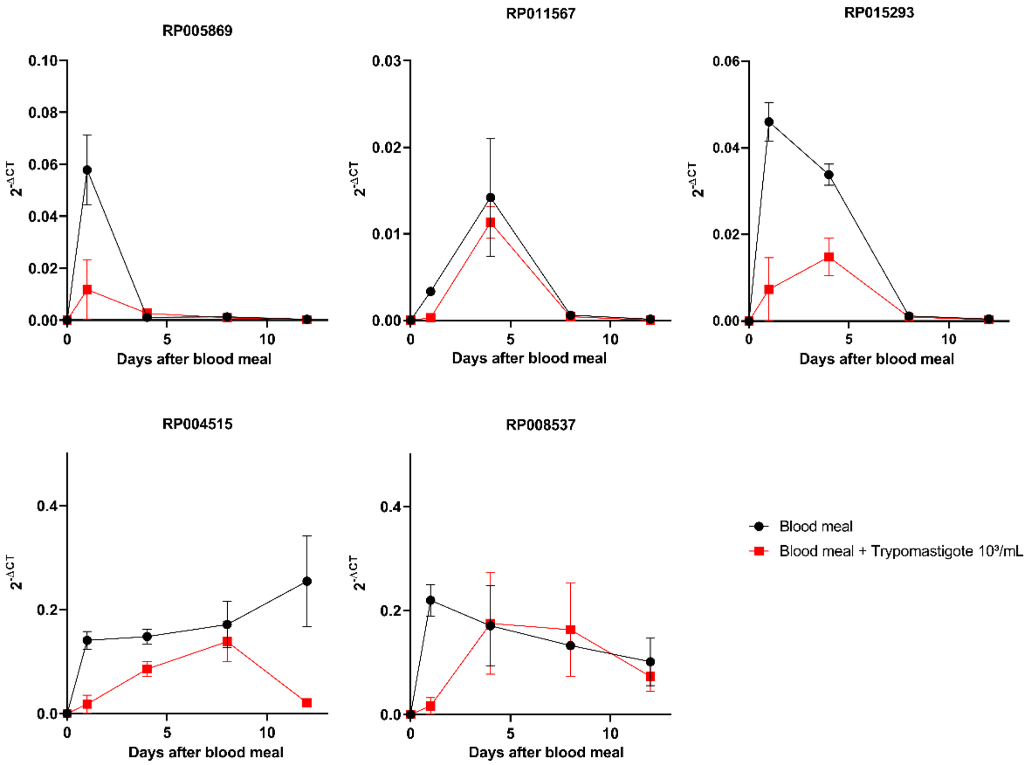


**Fig. S1.** qPCR evaluation of sHSPs expression profile after blood feeding (black dots) and after infection (CL-Brener clone, $10^3$ trypomastigote parasites/mL of blood; red squares). Experiments were performed under the same experimental design and conditions used for the transcriptome analysis. Data shown are mean ± SEM (n=5). Each sample is a pool of 10 midguts.

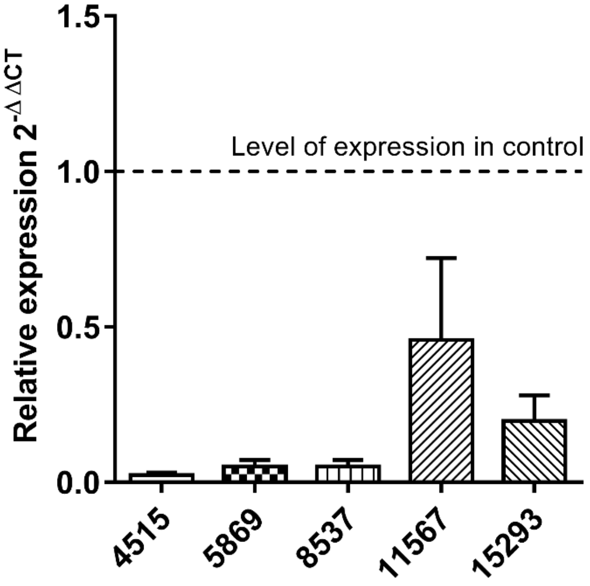


**Fig. S2.** *R. prolixus* females injected with dsRNA for sHSPs produce silenced offspring. Data shown are relative expression of each of the five sHSPs 4 days after blood meal of first instar nymphs hatched from eggs laid by females injected with a mix of the five dsRNAs hereafter referred to as dsMix. Nymphs were fed 5-7 days after hatching. Transcript levels were compared with control (dsMal; dotted line). Numbers under the X-axis refer to each sHSP transcript number.

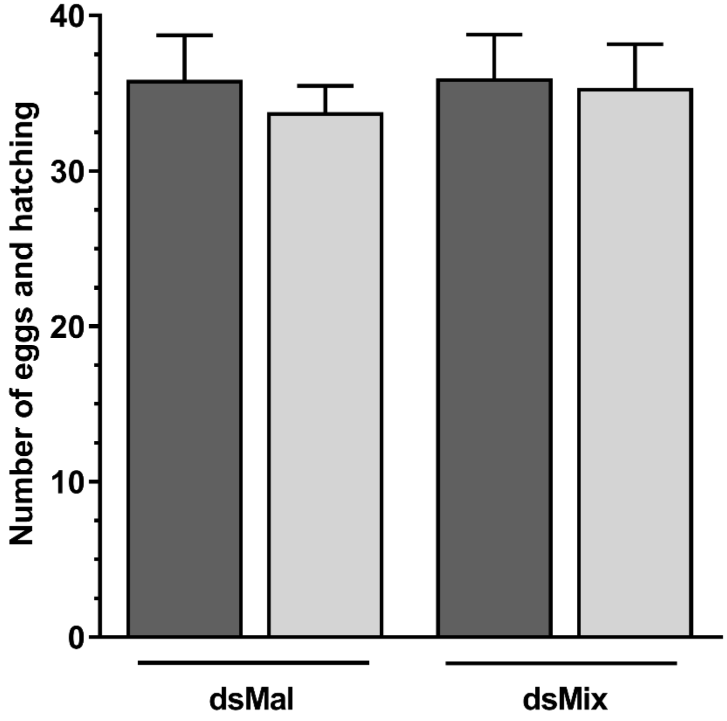


**Fig. S3.** Knockdown by dsMix injection into adult females did not affect oviposition (Dark grey) and hatching (light grey). Oviposition was quantified individually and plotted together. (n=12 females per point).

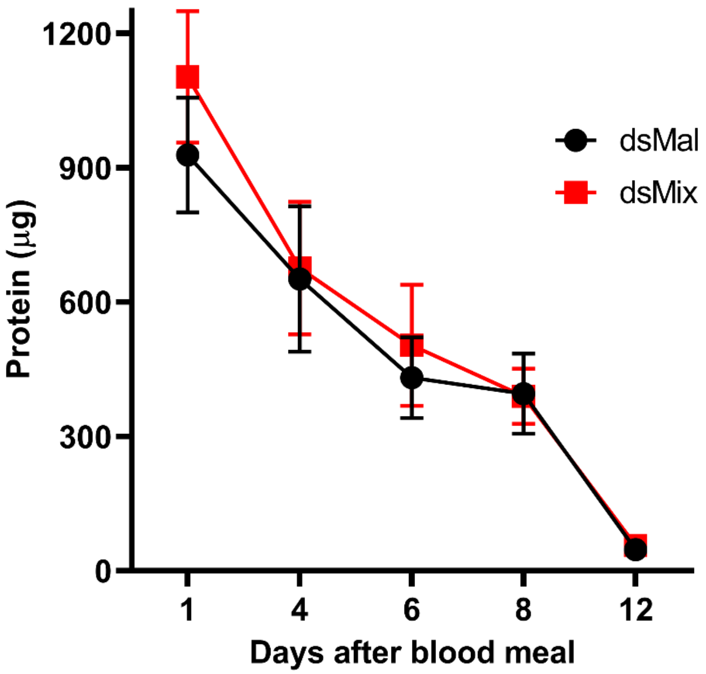


**Fig. S4.** Protein content of midgut from first instar nymphs of dsMal and dsMix insects was used to evaluate the time course of digestion of blood meal (ten first stage nymphs were individually assayed per day).

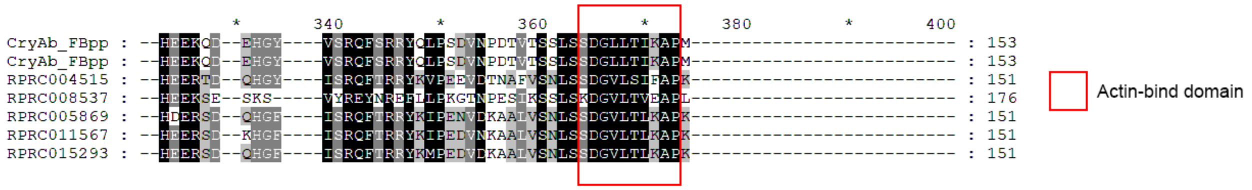


**Fig. S5.** Alignment of two *Drosophila* sHSPs sequences (FlyBase ID: FBgn0011296) at the carboxy terminal region of the protein (1) with the five *R. prolixus* sHSPs, showing the presence of a conserved actin-binding domain (red square) (2). Alignments were performed with SEAVIEW and the alignment image was processed using GeneDoc software.

**Table S1.** Oligonucleotide sequences for RT qPCR used in this study.

| Gene | Sense | Sequence 5'-3' |
|---|---|---|
| RP-004515 sHSP | Foward | ATTACAGACGCCCATTTCGT |
| | Reverse | TAATGCGGATACACCGGAAT |
| RP-005869 sHSP | Foward | GAACATGGCTGCTGGAGATT |
| | Reverse | GGGATCTTATAGCGGCGAGT |
| RP-008537 sHSP | Foward | GCCAATATCAACCCGAAGAA |
| | Reverse | AGTTTGTCTGGTCCCGTCAG |
| RP-011567 sHSP | Foward | CGGTTTCATCTCAAGGCAAT |
| | Reverse | GGAACTTCACGACCAGCTTT |
| RP-015293 sHSP | Foward | TCAGGGGTATCTGCGGTAAC |
| | Reverse | GTCCACATCCTCTGGCATTT |
| RP-EF-1 | Foward | GATTCCACTGAACCGCCTTA |
| | Reverse | GCCGGGTTATATCCGATTTT |
| Tarleton | Foward | CTCTTGCCCACAMGGGTGC |
| | Reverse | CCAAGCAGCGGATAGTTCAGG |
| Mal | Foward | AGCCCTCCCGTATCGTAGTT |
| | Reverse | CGATTCGGCCTATTGGTTA |

**Table S2.** Oligonucleotide sequences for dsRNA synthesis used in this study.

| Gene | Sense | Sequence 5'-3' |
| --- | --- | --- |
| **RP-004515 sHSP** | Foward | taatacgactcactatagggTCGGACAGAAGCGAATTTAGA |
| | Reverse | taatacgactcactatagggAAGGAAAAGTTTGATTTTTGTTCA |
| **RP-005869 sHSP** | Foward | taatacgactcactatagggCGTTCAACAGTTTAAGCCAGAA |
| | Reverse | taatacgactcactatagggTCTTCCTTCTTCTTTGCAGGA |
| **RP-008537 sHSP** | Foward | taatacgactcactatagggAAAAGTTCAACGTGGCTGGA |
| | Reverse | taatacgactcactatagggAGTTTGTCTGGTCCCGTCAG |
| **RP-011567 sHSP** | Foward | taatacgactcactatagggCTCATCAGACGGTGTCCTCA |
| | Reverse | taatacgactcactatagggTCCCTGGAGAAGGGCTAAAT |
| **RP-015293 sHSP** | Foward | taatacgactcactatagggGGATGTACAACAGTTCAAACCAGA |
| | Reverse | taatacgactcactatagggAAAAATTCATCAAAACAAGATTGG |
| **dsMal T7*** | TAATACGACTCACTATAGGG<br>* Plasmid Litmus 28i-mal (New England Biolabs) | |

**Movie S1. (separate file).** Control dsMal (Day 1). Representative video showing normal peristaltic contractions in one dsMal nymph. Contractions initiate at the anterior midgut and progress continuously to the posterior region without interruption. Videos were recorded for 5 minutes. Videos used came from 3-5 insects registered in at least three independent replicates of the experiment.

**Movie S2. (separate file).** dsMix (Day 1). Representative video of one dsMix nymph exhibiting altered peristaltic activity. Contractions initiate but fail to propagate continuously along the midgut, resulting in interrupted and incomplete waves. Videos were recorded for 5 minutes. Videos used came from 3-5 insects registered in at least three independent replicates of the experiment.

**Movie S3. (separate file).** Control insect uninfected (Day 1). Representative video showing normal peristaltic contractions in a control nymph. Contractions propagate smoothly from the anterior to the posterior midgut without interruption. Videos were recorded for 5 minutes. Videos used came from 3-5 insects registered in at least three independent replicates of the experiment.

**Movie S4. (separate file). I**nfected insects (Day 1). Representative video of one *Trypanosoma cruzi*–infected nymph showing markedly reduced gut motility, followed by one *Trypanosoma cruzi*–infected nymph showing altered peristaltic motility, similar to the behavior observed in the dsMix group. Several infected insects displayed little to no detectable peristaltic contractions during the 5-minute recording period. Videos used came from 3-5 insects registered in at least three independent replicates of the experiment.
